# Necrosulfonamide causes oxidation of PCM1 and impairs ciliogenesis and autophagy

**DOI:** 10.1101/2023.05.05.539527

**Authors:** Clotilde C.N. Renaud, Carolina Alves Nicolau, Clément Maghe, Kilian Trillet, Jane Jardine, Sophie Escot, Nicolas David, Julie Gavard, Nicolas Bidère

## Abstract

Centriolar satellites are high-order assemblies, scaffolded by the protein PCM1, that gravitate as particles around the centrosome and play pivotal roles in fundamental cellular processes notably ciliogenesis and autophagy. Despite stringent control mechanisms involving phosphorylation and ubiquitination, the landscape of post-translational modifications shaping these structures remains elusive. Here, we report that necrosulfonamide (NSA), a small molecule known for binding and inactivating the pivotal effector of cell death by necroptosis MLKL, intersects with centriolar satellites, ciliogenesis, and autophagy independently of MLKL. NSA functions as a potent redox cycler and triggers the oxidation and aggregation of PCM1 alongside select partners, while minimally impacting the overall distribution of centriolar satellites. Additionally, NSA-mediated ROS production disrupts ciliogenesis and leads to the accumulation of autophagy markers, partially alleviated by PCM1 deletion. Together, these results identify PCM1 as a redox sensor protein and provide new insights into the interplay between centriolar satellites and autophagy.

## INTRODUCTION

Centriolar satellites are small membrane-less electron-dense particles of 70-100 nm in diameter that shuttle onto microtubules and concentrate around the centrosome ^1–3^. These composite proteinaceous structures play a pivotal role in regulating the composition and function of the centrosome. Accordingly, 40% of the >600 proteins identified in the interactome of 22 satellite components are shared with centrosome ^4,5^. A large body of work now supports a role for centriolar satellites in the formation of primary cilia, a microtubule-based organelle that protrudes from the cell body, senses and integrates extracellular signals to regulate cell proliferation, polarity, nerve growth, differentiation, or tissue maintenance ^6,7^. Notably, mutations in genes encoding centriolar satellite components lead to defects in ciliogenesis and cause a variety of diseases and syndromes termed ciliopathies ^8^. Nonetheless, centriolar satellites also exert functions beyond centrosome homeostasis maintenance and ciliogenesis, such as microtubule organization, aggresome clearance, stress response, neurogenesis, and autophagy ^9–16^.

Centriolar satellites are scaffolded by the Pericentriolar Material 1(PCM1) protein ^17,18^. The depletion of PCM1 disassembles centriolar satellites, causes the degradation of several of their components, and prevents the formation of primary cilia ^9,19,20^. PCM1 also binds and protects the autophagy regulators GABARAP from proteasomal degradation, thereby regulating the maturation of autophagosome ^13,14,21^. PCM1 is a 2,024 amino acids protein tightly regulated by subcellular location and post-translational modifications (PTMs) ^2^. For instance, PCM1 is redistributed in the cytosol of the cells during mitosis or when microtubules are dismantled with the microtubule inhibitor nocodazole, and in cells exposed to cellular stresses such as UV radiation, heat shock, and transcription block ^17,22,23^. PCM1 is also differentially phosphorylated during the cell cycle. Accordingly, the phosphorylation of PCM1 by the polo-like kinase 4 (PLK4) on its serine residue 372 during the G1 phase is crucial for its dimerization and the maintenance of centriolar satellites ^24^. By contrast, the phosphorylation of PCM1 at T^703^ by the Cyclin-dependent kinase 1 (CDK1) during the transition from the G2 phase to mitosis leads to the dismantlement of centriolar satellites ^25^. Moreover, PCM1 is also marked by ubiquitin chains, catalyzed by the E3 ligase Mindbomb 1 (MIB1), and this leads to proteasomal degradation and, thereby, disruption of centriolar satellites ^19,20,22,26^. Lastly, PCM1, together with several other centriolar satellite components, is cleaved by caspase-3 during apoptosis ^27^. Nonetheless, whether PCM1 undergoes additional PTM is unknown.

In this work, we show that the small molecule necrosulfonamide (NSA), a known inhibitor of the human effector of cell death by necroptosis MLKL ^28^, induces the oxidation and aggregation of PCM1 and additional centriolar satellite components in an MLKL-independent manner. While NSA does not disrupt the overall architecture of centriolar satellites, it efficiently blocks primary cilia formation in human and mouse cell lines. Additionally, our findings indicate that NSA elevates the abundance of the autophagy markers p62 (SQSTM1) and GABARAPL1, with these effects partially dependent on PCM1. Collectively, these data designate centriolar satellites as redox sensor structures and offer a novel perspective on the regulation of the dynamic relationship between centriolar satellites and autophagy.

## RESULTS

### Necrosulfonamide triggers the oxidation of PCM1 independently of MLKL

To identify novel compounds prone to altering PCM1 status and/or abundance, we devised a screen by immunoblotting analysis of a panel of 35 broadly used small molecules targeting known signaling pathways in the human lymphoblastoid T cell line Jurkat (Table S1). The best hit was necrosulfonamide (NSA), a small compound previously shown to bind and inhibit the human pseudokinase MLKL, a crucial effector of cell death by necroptosis and extracellular vesicle biogenesis ^28–30^. When applied at non-toxic concentrations, NSA led to the appearance of slower migration species of PCM1 combined with a decrease of full-length PCM1 (Fig. 1A and 1B). Of note, we did not observe significant changes in the cell cycle in Jurkat cells treated with NSA (Fig. S1A). A similar alteration in the PCM1 profile was observed in the retinal-pigmented epithelial (RPE-1) cell line, indicating that this effect was not restricted to a particular cell lineage (Fig. S1B). At the mRNA level, no overt change in PCM1 and other centriolar satellite components, except that of CEP131, was observed in NSA-treated Jurkat cells, suggesting PTM (Fig. S1C).

**Figure 1.**
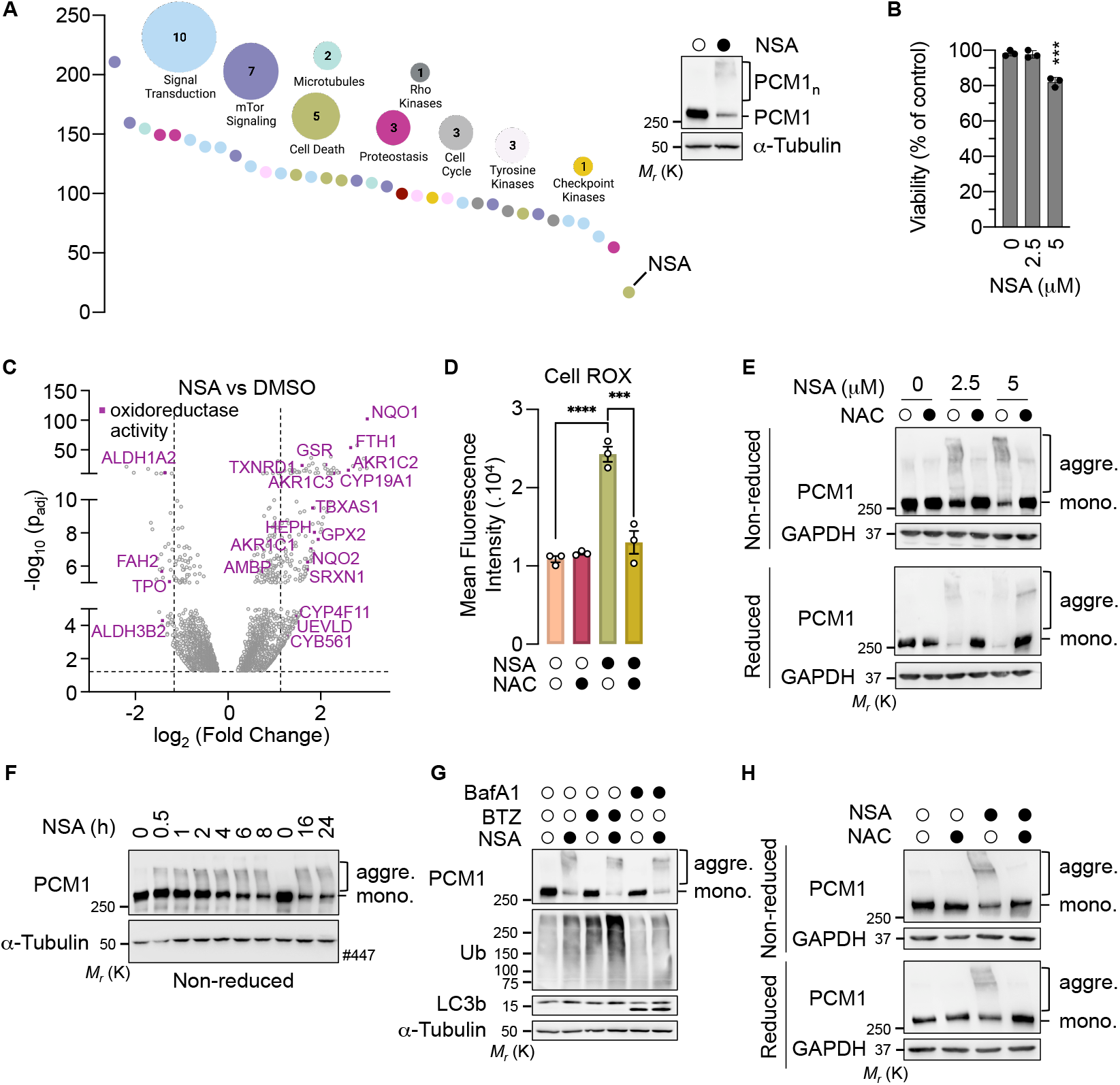
Necrosulfonamide causes the oxidation and aggregation of PCM1 independently of MLKL (A) Densitometric measurement of PCM1 abundance, normalized to GAPDH and compared to DMSO-treated samples analyzed by Western blotting in Jurkat cells treated overnight with 35 small-molecule compounds. The inset blot shows samples from cells treated with NSA (2.5 μM), or with DMSO overnight analyzed by Western blotting as indicated. Molecular weight markers (*Mr*) are indicated. PCM1_n_ indicates high-molecular-weight species of PCM1. **(B)** CellTiter-Glo Luminescent cell viability assay of Jurkat cells treated overnight with 2.5 or 5 μM NSA (biological triplicates, ^***^*P* < 0.001, ANOVA). **(C)** RNAseq transcriptomic analysis of Jurkat cells treated with DMSO or 2.5 μM NSA. The scatter (volcano) plot shows Log2 Fold Change and -Log10 (padj); n=3. Dashed lines, significance cut-off. Purple symbols indicate differentially expressed genes associated with the main function “oxidoreductase activity”. **(D)** Flow cytometry analysis of ROS using CellROX in Jurkat cells pretreated with 5 mM N-acetylcysteine (NAC) for 2h and exposed to 2.5 μM NSA overnight (mean ± SEM; n=3; ^***^*P* < 0.001, ^****^*P* < 0.0001; ANOVA). **(E)** Jurkat cells were pretreated with 5 mM NAC for 2h before overnight incubation with NSA, as indicated. Cell lysates were analyzed by Western blotting in reducing (10% 2β-ME) and non-reducing conditions. Aggre., aggregates; mono., monomeric. **(F)** Jurkat cells were treated with 2.5 μM NSA or DMSO, as indicated. Cell lysates were analyzed by Western blotting in non-reducing conditions. **(G)** Jurkat cells were pretreated with 10 nM Bortezomib (BTZ) or 100 nM Bafilomycin A1 (BafA1) for 1h and incubated with 2.5 μM NSA overnight. Cell lysates were analyzed by Western blotting with antibodies specific to the indicated proteins. **(H)** L929 cells were treated and analyzed as in (E). All presented data are representative of three independent experiments.

To gain insights into the signaling pathways that could lead to PCM1 alteration, we next conducted an RNA-sequencing analysis of Jurkat cells treated overnight with NSA. This identified 3,648 differentially expressed genes, out of which 235 (39 down and 196 up) were enriched with a shrunken Log2 fold change of at least 1.2. The term “oxidoreductase activity” emerged among the top differentially regulated pathways, suggesting a link between NSA and the production of reactive oxygen species (ROS) (Fig. 1C). Moreover, this dataset unveiled a signature for the NRF2 (also named NFE2L2), a master transcriptional controller of antioxidant genes ^31^ (Fig. S1D). Accordingly, NSA treatment resulted in increased expression of mRNA encoding for the NRF2 targets NQO1 and GSR. This upregulation was suppressed in cells pretreated with the anti-oxidant N-acetylcysteine (NAC) (Fig. S1E). Of note, the silencing of PCM1 did not drastically impact NSA-mediated induction of these genes (Fig. S1E). In line with this, the analysis of ROS production by flow cytometry showed a significant increase in live cells exposed to NSA, which was hampered by pretreatment with NAC (Fig. 1D). Interestingly, NSA also led to a NAC-dependent augmentation in green fluorescence (Fig. S1F). This may reflect an increase in the abundance of the ROS detoxification flavoproteins, which are major contributors to autofluorescence ^32^. Oxidative stressors can cause covalent oligomerization of proteins as well as the formation of disulfide-linked conjugates ^33–35^. We therefore hypothesized that the high-order species of PCM1 may be aggregates that resist our reducing conditions. Accordingly, the analysis by immunoblotting of cell lysates in reducing and non-reducing conditions showed that NAC efficiently prevented the formation of highmolecular-weight species of PCM1 (Fig. 1E). By contrast, MLKL remained monomeric in all conditions tested, indicating that it is likely not oxidated (Fig. S1G). We further observed that PCM1 aggregates formed rapidly following NSA treatment and remained stable after several hours of treatment (Fig. 1F). Moreover, the blockage of proteasomal and lysosomal degradation pathways with Bortezomib and Bafilomycin A1 (BafA1), respectively, did not overtly change the pattern of PCM1, suggesting that PCM1 aggregates are stable structures and likely not prone to degradation (Fig. 1G). Collectively, these results suggest that NSA is a redox cycler that drives the aggregation of PCM1.

NSA binds the Cysteine in position 86 of human MLKL, inhibiting its oligomerization and subsequent programmed cell death by necroptosis ^28^. To further explore whether MLKL participates in PCM1 aggregation, its expression was targeted with small interfering RNA (siRNA). However, high molecular weight species of PCM1 normally appeared in MLKL-silenced cells treated with NSA (Fig. S1H). In mouse MLKL, the Cys^86^ is substituted with a Tryptophan residue, making NSA inefficient ^28^. Nevertheless, we observed that NSA efficiently drove PCM1 aggregation in the mouse cell line L929 and mouse embryonic fibroblasts (MEFs) (Fig. 1H, and S1I), further supporting an MLKL-independent action. Altogether, this suggests that NSA drives PCM1 aggregation independently of MLKL.

### Necrosulfonamide inhibits the formation of primary cilia

We next explored the impact of NSA on the distribution of centriolar satellites. Confocal microscopy analysis showed that the location of PCM1 around the centrosome was not overtly changed in NSA-treated cells (Fig. 2A and 2B). The same was true in RPE-1 cells (Fig. S2A). Moreover, the fluorescence intensity of PCM1 staining in the vicinity of the centrosome did not decrease with NSA, further arguing against a global degradation of PCM1 (Fig. 2C). We further assessed whether NSA alters additional components of centriolar satellites and the centrosome by immunoblotting. We found that some proteins such as CEP131, CEP72, OFD1, and to a lesser extent CEP290 formed ROS-dependent aggregates, whereas others including MIB1 and γ-Tubulin were essentially monomeric. Of note, the abundance of MIB1, which is elevated in the absence of PCM1 ^19,20^, remained unchanged in cells treated with NSA (Fig. 2D). In addition, PCM1 efficiently co-immunoprecipitated with its known partners CEP290, MIB1, and CEP131 in the presence of NSA (Fig. 2E). Hence, these data suggest that NSA induces the aggregation of PCM1 and several centriolar satellite components but does not compromise its ability to interact with its partners or its spatial organization in cells.

**Figure 2.**
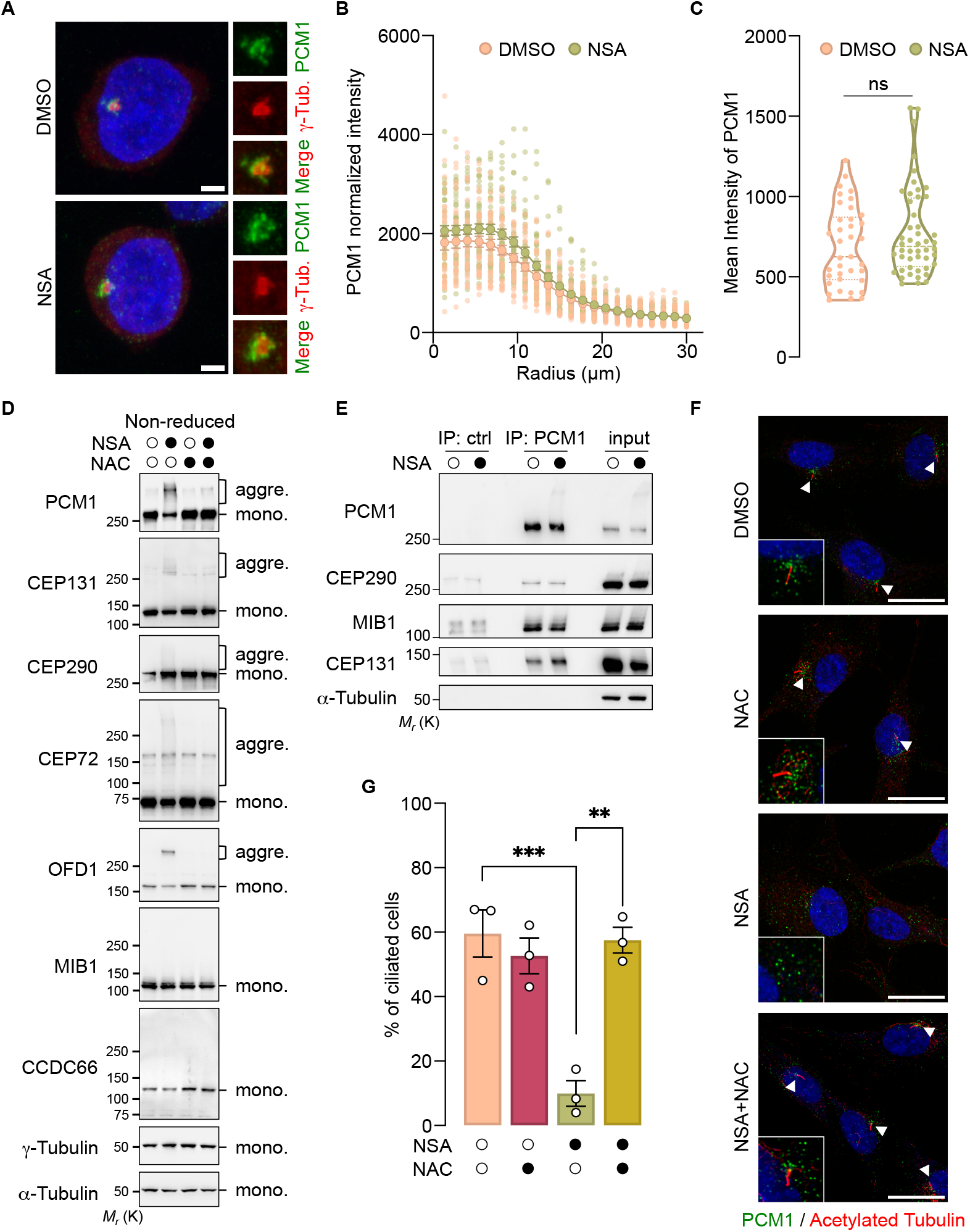
Necrosulfonamide inhibits ciliogenesis. (**A-C**) Confocal microscopy analysis of Jurkat cells treated overnight with 2.5 μM NSA showing the localization of PCM1 and γ-Tubulin, with nuclei stained by 4’-6-diamidino-2-phenylindole (DAPI). Scale bar, 2 μm. The radial profile (B) and the mean intensity (C) of PCM1 around the centrosome defined by γ-Tubulin were measured (n=37 (DMSO) and n=48 (NSA) cells from of three independent experiments; ns, non-significant; ANOVA). **(D)** Jurkat cells were pretreated with 5 mM N-acetylcysteine (NAC) for 2h and incubated with 2.5 μM NSA overnight. Cell lysates prepared in non-reduced conditions were analyzed by Western blotting with antibodies specific to the indicated proteins. Molecular weight markers (*Mr*) are indicated. Aggre., aggregates; mono., monomeric. **(E)** Cell lysates from Jurkat cells treated with 2.5 μM NSA overnight were subjected to immunoprecipitation (IP) with antibodies against PCM1 or with control (ctrl) antibodies, and samples were then analyzed by Western blotting as indicated. (**F and G**) RPE-1 cells were serum starved and treated with 2.5 μM NSA with and without NAC (5 mM) for 24 h and were then analyzed by confocal microscopy to visualize PCM1 and primary cilia (acetylated tubulin, arrowheads). Scale bar, 20 µm. In (F), the histogram shows the quantification of cells with cilia (mean ± SEM of three independent experiments; n > 100 cells counted per sample; ^**^*P* < 0.01, ^***^*P* < 0.001, ANOVA). All presented data are representative of three independent experiments.

Because PCM1 is instrumental for ciliogenesis ^9^, we next assessed the influence of NSA on the formation of primary cilia. To this end, RPE-1 cells were serum-starved to induce ciliogenesis, and acetylated tubulin, a constituent of centriole and the primary cilium, was tracked by confocal microscopy. We observed that treatment with NSA led to a significant reduction in the number of cells harboring an elongated acetylated tubulin staining, suggesting impaired ciliogenesis (Fig. 2F and 2G). However, this defect in ciliogenesis was overcome by pretreating the cells with NAC (Fig. 2F and 2G). Of note, MLKL silencing had no overt impact on ciliogenesis induced by serum starvation or on the ability of NSA to prevent it, suggesting an MLKL-independent function (Fig. S2B). In line with this, NSA hampered serum starvation-induced ciliogenesis in MEFs, in which NSA cannot bind MLKL (Fig. S2C). Altogether, this suggests that NSA-mediated ROS production alters PCM1 and ciliogenesis in an MLKL-independent manner.

### Necrosulfonamide leads to defects in autophagy

In addition to oxidative stress, our transcriptomic analysis identified a signature related to endosomes and lysosomes in NSA-treated samples (Fig. 3A). Moreover, the screen of a qPCR-based array of autophagy regulators also indicated an up-regulation of some autophagy actors, including GABARAPL1, p62, and LC3 in NSA-treated cells (Fig. 3B). Accordingly, we observed a significant increase in GABARAPL1 and p62 protein levels in response to NSA treatment by immunoblotting analyses (Fig. 3C). p62 is a well-established redox sensor ^33–35^, and we found that NSA drove its oxidation. This was however not the case for GABARAPL1. Importantly, the upregulation of GABARAPL1 and p62, as well as p62 oxidation were ROS-dependent (Fig. 3C). The NSA-dependent increase in p62 and GABARAPL1 abundance was also seen in mouse cells, suggesting again a role of NSA independent of MLKL (Fig. S3A and S3B). Nonetheless, this increase in p62 and GABARAPL1 abundance in NSA-treated cells could reflect an augmented transcription and/or a decreased degradation. We therefore assessed the impact of NSA on the autophagic flux. To this end, Jurkat cells were treated with the V-ATPase inhibitor Bafilomycin A1 (BafA1), which blocks autophagic degradation. Our immunoblotting analysis showed that the accumulation of p62 and GABARAPL1 driven by NSA was further increased with BafA1, suggesting a transcription-dependent increase in autophagy markers (Fig. S3C). A similar pattern was observed in RPE-1 cells (Fig. S3D). In addition, the analysis of RPE-1 cells by confocal microscopy unveiled a more punctuate staining of p62 and GABARAPL1 in the presence of NSA, which was further increased with BafA1 (Fig. 3D-3G). Of note, NSA-mediated upregulation of GABARAPL1 was abolished in cells treated with the protein synthesis inhibitor cycloheximide (Fig. S3E). Collectively, our data suggest that NSA increases the abundance of some of the autophagy markers at RNA and protein levels.

**Figure 3.**
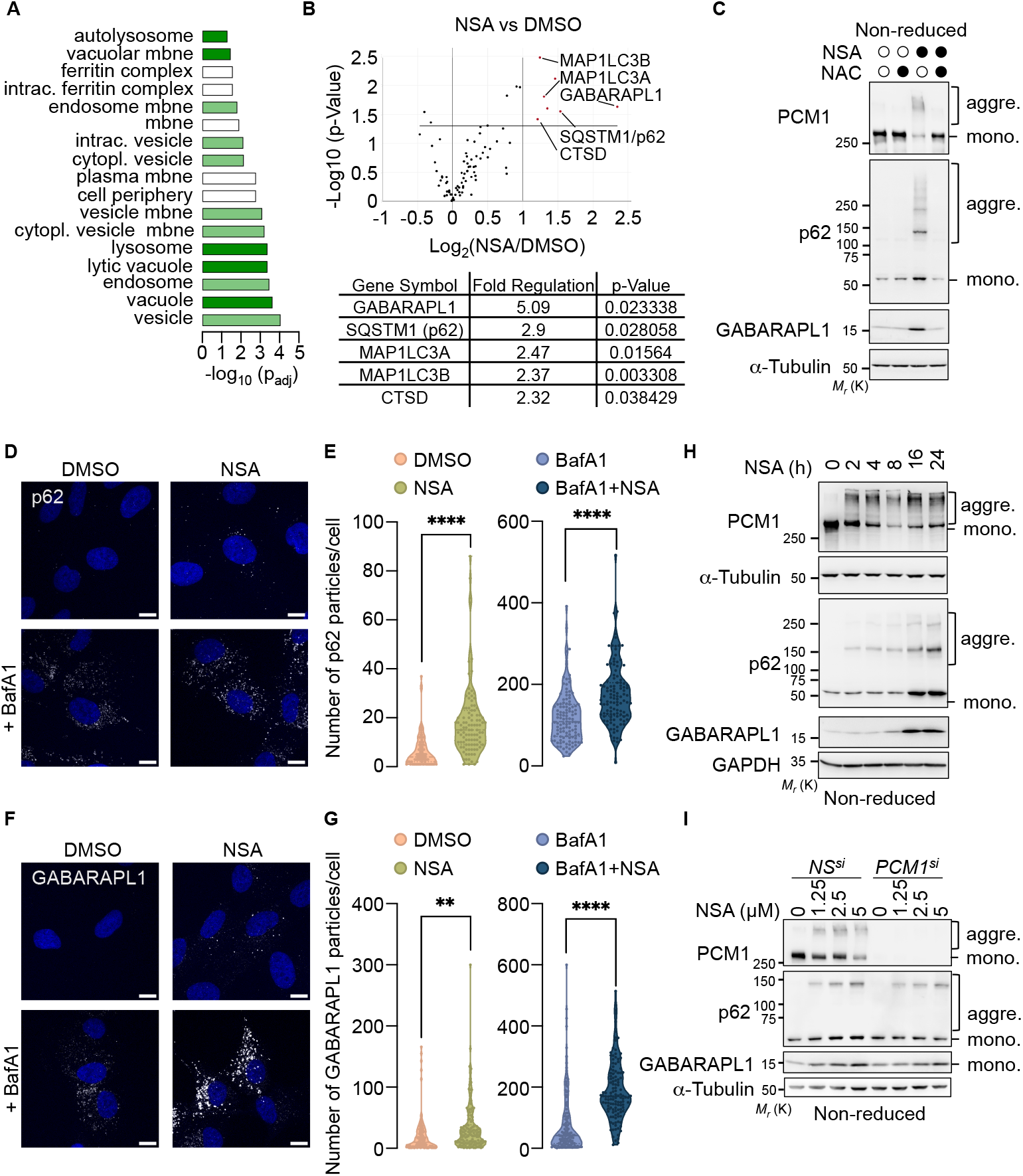
Necrosulfonamide causes a defect in autophagy. **(A)** Cellular Component enrichment analysis of differentially expressed genes analyzed from an RNA-seq analysis of Jurkat cells treated overnight with 2.5 μM NSA, as described in Fig. 1C. **(B)** Volcano plot of RT^2^ profiler PCR array of human autophagy signaling components for Jurkat cells treated as in (A). Data represent three independent experiments. Genes up-regulated upon NSA treatment are shown. **(C)** Jurkat cells pre-incubated with 5 mM N-acetylcysteine (NAC) were treated with 2.5 μM NSA overnight. Cell lysates prepared in non-reduced conditions were analyzed by Western blotting with antibodies specific to the indicated proteins. Aggre., aggregated proteins; mono., monomeric proteins. Molecular weight markers (*Mr*) are indicated. (**D and E**) Confocal microscopy analysis of p62 in RPE-1 cells pretreated with 100 nM Bafilomycin A1 (BafA1) for 1h and incubated with 2.5 μM NSA overnight. Nuclei were stained by DAPI. Scale bar, 10 µm. The number of p62 puncta per cell is shown in (E) (n>96 cells analyzed per sample of four independent experiments, ^****^*P* < 0.0001, t-test). (**F and G**) Confocal microscopy analysis of GABARAPL1 in RPE-1 cells as in (D and E) (n>113 cells analyzed per sample of four independent experiments, ^**^*P* < 0.01, ^****^*P* < 0.0001, t-test). (**H**) Cell lysates from Jurkat cells treated with 2.5 μM of NSA as indicated were prepared in non-reduced conditions and analyzed by Western blotting with antibodies against the indicated proteins. (**I**) Jurkat cells were transfected with small interfering RNA (siRNA) for PCM1 or scramble non-specific (NS). Cells were treated overnight with NSA as indicated. Cell lysates were prepared and analyzed by Western blotting with antibodies specific against the indicated proteins. All presented data are representative of three independent experiments.

Lastly, we wondered whether NSA drives PCM1 aggregation and alteration of autophagy markers in a linear or parallel manner. First, a time-course experiment by immunoblotting showed that NSA-mediated PCM1 aggregation preceded the accumulation of p62 and GABARAPL1 (Fig. 3H). PCM1 was then silenced with siRNA in Jurkat cells. At the mRNA level, we found that although PCM1 silencing elevated the transcription of GABARAPL1, it did not alter the upregulated transcription of GABARAPL1, p62, and LC3 induced by NSA (Fig. S3F). As expected, treating cells with the anti-oxidant NAC counteracted NSA-mediated induction of autophagy markers (Fig. S3F). Nonetheless, our immunoblot analysis showed that the increase in the abundance of GABARAPL1 caused by NSA was diminished without PCM1 (Fig. 3I). Similarly, PCM1-silenced cells exhibited reduced NSA-mediated p62 upregulation. Taken together, these results suggest that the accumulation of autophagy markers induced by NSA treatment partially depends on PCM1 at the post-transcriptional level.

## DISCUSSION

NSA is an alkylating agent targeting cysteine residues, that was initially found to bind to the Cys^86^ of human MLKL ^28^. NSA disrupts disulfide bonds formed by MLKL and thereby prevents MLKL oligomerization and insertion in the plasma membrane, a crucial step in the execution of cell death by necroptosis ^28^. Recent studies have also ascribed a role for NSA in limiting cell death by pyroptosis ^36,37^, although the underlying mechanism of action is still debated ^36,37^. However, the full spectrum of NSA’s targets remains unclear. We now report that NSA acts as a potent redox cycler, impacting ciliogenesis and autophagy via ROS independently of MLKL. These results support the recent description that NSA drives ROS generation in monocytic cell lines ^38^. Our data demonstrate rapid aggregation of PCM1, several centriolar satellite components, and the established redox protein p62 upon NSA exposure. Nonetheless, further investigation is necessary to delineate the global impact of NSA on the transcriptome and proteome of the cells and to identify the landscape of its direct and indirect targets. Additionally, this function we ascribe to NSA, distinct from its established inhibitory role on MLKL, may hold implications for the field of cell death by necroptosis.

PCM1, the linchpin for centriolar satellites, undergoes tight regulation via phosphorylation, ubiquitination, and proteolysis ^2,27^. Our study now expands the landscape of PCM1 PTMs to oxidation and aggregation. Furthermore, besides PCM1, we observed several other centriolar components acting as redox proteins. Nonetheless, we found that centriolar satellites normally orbit the centrosome when their components form aggregates. This aligns with previous research indicating that the ROS H_2_O_2_ does not disrupt the spatial distribution of centriolar satellites ^39^. Our data show that NSA prevents the formation of primary cilia, a process controlled by centriolar satellites. However, the contribution of centriolar satellite oxidation remains to be evaluated. We also found that NSA had a large influence on autophagy, acting both at the transcriptional and post-transcriptional levels. First, we observed that NSA drives the transcription of pivotal autophagy markers like p62 and GABARAPL1 via ROS induction. Our transcriptomic dataset was associated with the activation of NRF2, a crucial transcriptional regulator of antioxidant genes ^31^. It is tempting to speculate that NRF2 is activated following NSA treatment, as it has been shown to stimulate the expression of p62 and GABARAPL1 ^31^. Additionally, NSA drives the oxidation and aggregation of p62 through ROS generation, a critical step in p62-mediated pro-survival autophagy in stress conditions ^33–35^. Importantly, we found that NSA-mediated upregulation of p62 and GABARAPL1 partly requires centriolar satellites at the post-transcriptional level, as evidenced in PCM1-silenced cells. Yet, defining whether this involves the oxidation of centriolar satellite components will require additional work. In addition, the oxidation of centriolar satellites may regulate different stages of the autophagic process. For instance, PCM1 interacts with GABARAPL1 via an LC3-interacting region (LIR) motif and finely tunes autophagy ^13,14^, whereas OFD1 negatively regulates autophagosome biogenesis ^40^. Lastly, autophagy has been proposed to control the abundance of ciliary proteins, including IFT20 and OFD1 ^11,12^, and it would be of interest to evaluate how NSA-mediated dysregulation of autophagy intersects with ciliogenesis. In addition to primary cilia formation and autophagy, centriolar satellites play a vital role in various cellular functions, such as the clearance of pathogenic aggregates containing misfolded proteins, microtubule organization, and neurogenesis ^2^. Hence, our future work will be aimed at characterizing how oxidation shapes the functions of centriolar satellites.

## Limitations of the study

Our study indicates that NSA displays a redox cycler activity independent of its described target MLKL. We conclusively showed that NSA generates ROS, which cause the aggregation of PCM1 and other centriolar satellite components, and alter key cellular functions such as ciliogenesis and autophagy. Although PCM1 orchestrates the formation of primary cilia and has been associated with autophagy, we have not formally demonstrated the link between these observations and PCM1 oxidation. Future studies will therefore be required to identify the oxidized residues and the impact of their modification on the functions controlled by PCM1.

## ACKNOWLEDGMENTS

The authors would like to thank the MicroPicell facility (SFR Santé François Bonamy, Nantes, France) for the expertise and discussions for imaging analysis. This work was funded by Fondation de France, Fondation ARC contre le Cancer (PJA to J.G. and N.B.), Institut National du Cancer (INCa PAIR-CEREB lNCa_16285, INCa_18384), Ligue nationale contre le cancer (Equipe labellisée) et comités de Loire-Atlantique, Maine et Loire, Vendée, Ille-et-Vilaine, Mayenne, and Finistère (J.G., N.B.). C.C.N.R. received a fellowship from INSERM, Région Pays de la Loire, and the Ligue Nationale contre le Cancer. C.A.N. and J.J. were supported by Fondation de France and C.M. by Ligue Régionale contre le Cancer and Région Pays-de-la-Loire. The team is part of the SIRIC ILIAD (INCA-DGOS-INSERM-ITMO Cancer_18011).

## AUTHOR CONTRIBUTIONS

Conceptualization, C.C.N.R., C.A.N., C.M., K.T., J.J., S.E., N.D., J.G., and N.B.; Methodology, C.C.N.R., C.A.N., C.M., K.T., J.J., and N.B.; Investigation, C.C.N.R., C.A.N., C.M., K.T., J.J., and N.B.; Supervision: N.B.; Writing-Original Draft, C.C.N.R., C.A.N., and N.B.; Writing-Review and Editing, C.C.N.R., C.A.N., C.M., K.T., J.J., J.G., and N.B.; Funding Acquisition, J.G., and N.B.

## METHODS

This study does not involve any patients or healthy control participants.

### Cultured Cells

Human hTERT RPE-1, Jurkat E6.1, and L929 cells were purchased from the American Type Culture Collection (ATCC). hTERT RPE-1 cells were cultured in Dulbecco’s modified Eagle’s F12 (DMEM: F12, Life Technologies) supplemented with 10 % heat-inactivated fetal bovine serum (FBS), glutamine, and Penicillin/Streptomycin (Life Technologies). Jurkat E6.1 cells were maintained with RPMI1640 (Life Technologies) supplemented with 10% FBS, HEPES (Life Technologies), Sodium Pyruvate (Life Technologies), and Penicillin/Streptomycin. MEFs and L929 cells were grown in Dulbecco’s modified Eagle’s (DMEM, Life Technologies) supplemented with 10% FBS, glutamine, and Penicillin/Streptomycin. Ciliogenesis was induced by washing and incubating RPE-1 or MEFs with OPTI-MEM (Life Technologies) for 24h.

### Reagents and Chemical Screen

The list of chemicals used in this work is provided in the key resources table. For the chemical screening, Jurkat cells were treated overnight with vehicle (DMSO) and 35 approved drugs (Table S1) targeting cell death, microtubules, mTOR signaling pathway, proteostasis, antigen receptor signaling, checkpoint kinases, cell cycle, tyrosine-kinase, and Rho-kinases. Cell lysates were prepared and analyzed by Western blot, as described below. Cell viability was assessed by CellTiter-Glo (Promega) following the manufacturer’s instructions. The cell cycle was analyzed using a NucleoCounter NC-300 fluorescent imaging cytometer as per the manufacturer’s instructions.

### siRNA Transfection

RPE-1 cells were transfected with 10 pmol of individual siRNA, at a final concentration of 16.5 nM, using the Lipofectamine RNAiMAX Transfection Reagent (Life Technologies), according to the manufacturer’s instructions. Jurkat cells were transfected by electroporation (BTX ECM 830, Harvard Apparatus) as previously described ^41^. The following sequences (Invitrogen, Stealth) were used: PCM1, GCCUAACCCUUUGCCGUUACGUUUA (HSS107661); MLKL, UCGAAUCUCCCAACAUCCUGCGUAU (HSS136795).

### Western blotting and Immunoprecipitations

Cells were washed with ice-cold PBS and pelleted by centrifugation at 300*g* before cell lysis with RIPA buffer [25 mM Tris-HCl (pH 7.4), 150 mM NaCl, 0.1% SDS, 0.5% Na-Deoxycholate, 1% NP-40, 1 mM EDTA] supplemented with protease inhibitors (Thermo Fisher Scientific) for 30 minutes on ice. Samples were cleared by centrifugation at 10,000*g* and protein concentration was determined by BCA (UP40840A; Interchim). 5-10 μg proteins in reducing (10% 2β-ME) and non-reducing conditions were resolved by SDS-PAGE using either 5-20% Tris-glycine or 3-15% Tris-acetate gels. The proteins were then transferred to nitrocellulose membranes (GE Healthcare). For immunoprecipitations, samples were pre-cleared with Protein G Sepharose (GE Healthcare) for 1h and subsequently incubated with 1-2 μg of antibodies for 2h at 4°C. The beads were pelleted by centrifugation at 5,000*g*, washed four times with lysis buffer, and eluted with 2x Laemmli sample buffer at 95°C for 5 min before resolving by SDS-PAGE.

Antibodies specific for the following proteins were purchased from Santa Cruz Biotechnology: PCM1 (G-6), Acetylated Tubulin (6-11 B-1), GAPDH (6C5), α-Tubulin (TU-02), γ-Tubulin (TU-30), Ubiquitin (P4D1), MIB1 (B-9), CEP290 (B-7). Antibodies specific to PCM1 (G2000), MLKL (D2I6N), SQSTM1/p62 (D5L7G), and SQSTM1/p62, LC3B (D11) were purchased from Cell signaling. Antibodies against GABARAPL1 (11010-1-AP) and α-Tubulin (66031-1-1g) were from Proteintech. Antibodies to CEP131 (A301-415A) and CCDC66 (A303-339A) were from Bethyl Laboratories. Antibodies against OFD1 (HPA031103) and PCM1 (ab72443) were purchased from Atlas and Abcam, respectively.

### Reactive Oxygen Species (ROS) Detection

Jurkat cells were pretreated with 5 mM N-Acetyl-L-Cysteine (NAC) for 2h before incubation overnight with 2.5 μM NSA. ROS levels were detected using CellROX (C10422, Life Technologies) by flow cytometry. Flow cytometry analyses were performed on FACS Calibur (BD Biosciences; Cytocell Facility, SFR François Bonamy, France) and processed using FlowJo V10 software.

### RNAseq

Jurkat cells were treated with vehicle (DMSO) or NSA (2.5 μM) overnight, in three independent experiments, washed with PBS, and snap-frozen on dry ice. RNA extraction (all RNA integrity number > 9.0), library preparation, RNAseq, and bioinformatics analysis were performed at Active Motif (Carlsbad, California, USA). Briefly, 2 µg of total RNA isolated using the NucleoSpin RNA plus Mini Kit for RNA Purification were further sequenced in Illumina sequencing (using NextSeq 500). The paired-end 42 bp sequencing reads (PE42) generated by Illumina were mapped to the genome using the STAR algorithm - DESeq2 software pipeline described in the Data Explanation document. Genes considered differently expressed had an adjusted p-value of less than 0.1. Enrichment analysis of the RNAseq experiment were performed using g:Profiler (version e107_eg54_p17_bf42210) applying a significance threshold of 0.01 and a |log_2_(fold change)|>1.2.

### qPCR and Autophagy Array

1.10^6^ RPE-1 cells or 4.10^6^ Jurkat cells were treated with vehicle (DMSO), NAC (5mM for 2h) and NSA (2.5 μM) overnight in biological triplicates. The cells were harvested in PBS, snapfrozen, and the RNA extraction was done using a NucleoSpin RNA plus Mini Kit for RNA Purification. Equal amounts of RNA were reverse-transcribed using the Maxima Reverse Transcriptase kit, and 50 ng of the resulting cDNA was amplified by qPCR using PerfeCTa SYBR Green FastMix Low ROX (Quantabio) (Table S2). Data were analyzed using the 2-ΔΔCt methods and normalized by the housekeeping genes ACTB and HPRT1. See Table S2 for the complete list of primers used. The autophagy screening was performed using the RT2 Profiler™ PCR Array Human Autophagy (Qiagen) following the manufacturer’s instructions.

### Immunofluorescence

RPE-1 cells were seeded onto 12 mm coverslips (Life Technologies) and Jurkat cells were adhered on polysine slides for 15 min (VWR, 631-0107). For ciliogenesis and centriolar satellite staining, the cells were fixed in 4% paraformaldehyde for 12 min at room temperature, incubated with ice-cold methanol for 2 min, and quenched for 10 min with PBS/100 mM glycine. Fixed cells were incubated for 60 minutes with primary antibodies and 60 minutes with secondary antibodies in PBS 1X-0.2% BSA-0.05% Saponin. Coverslips were sealed with Prolong gold anti-fade mounting solution (Life Technologies), and nuclei were counterstained with 4’,6-diamidino-2-phenylindole (DAPI). The following primary antibodies were used: PCM1 (Cell signaling, 5213), γTubulin (Sigma, GTU88), and acetylated Tubulin (Santa Cruz Biotechnology, 6-11 B-1). To stain autophagy markers, cells were fixed in PBS-paraformaldehyde 4% and permeabilized with PBS 4%BSA 0.3% Triton X-100. The following primary antibodies were used: SQSTM1/p62 (Cell signaling, 88588) and GABARAPL1 (Proteintech, 11010-1-AP). Images were acquired on a Nikon A1 Rsi, using a 60× oil-immersion lens (Nikon Excellence Center, MicroPicell, SFR Francois Bonamy, Nantes, France). Images were processed using FIJI software.

## QUANTIFICATION AND STATISTICAL ANALYSIS

Particles quantification of p62 and GABARAPL1 was carried out on thresholded images and using the analyze particles plugin of FIJI software. Statistical analyses were performed using GraphPad Prism 8 (GraphPad Software) using Student t-test and one-or two-way analysis of variance (ANOVA). The significance and number of repeats are indicated in figure legends.

