## Supplemental files for "Necrosulfonamide causes oxidation of PCM1 and impairs ciliogenesis and autophagy"

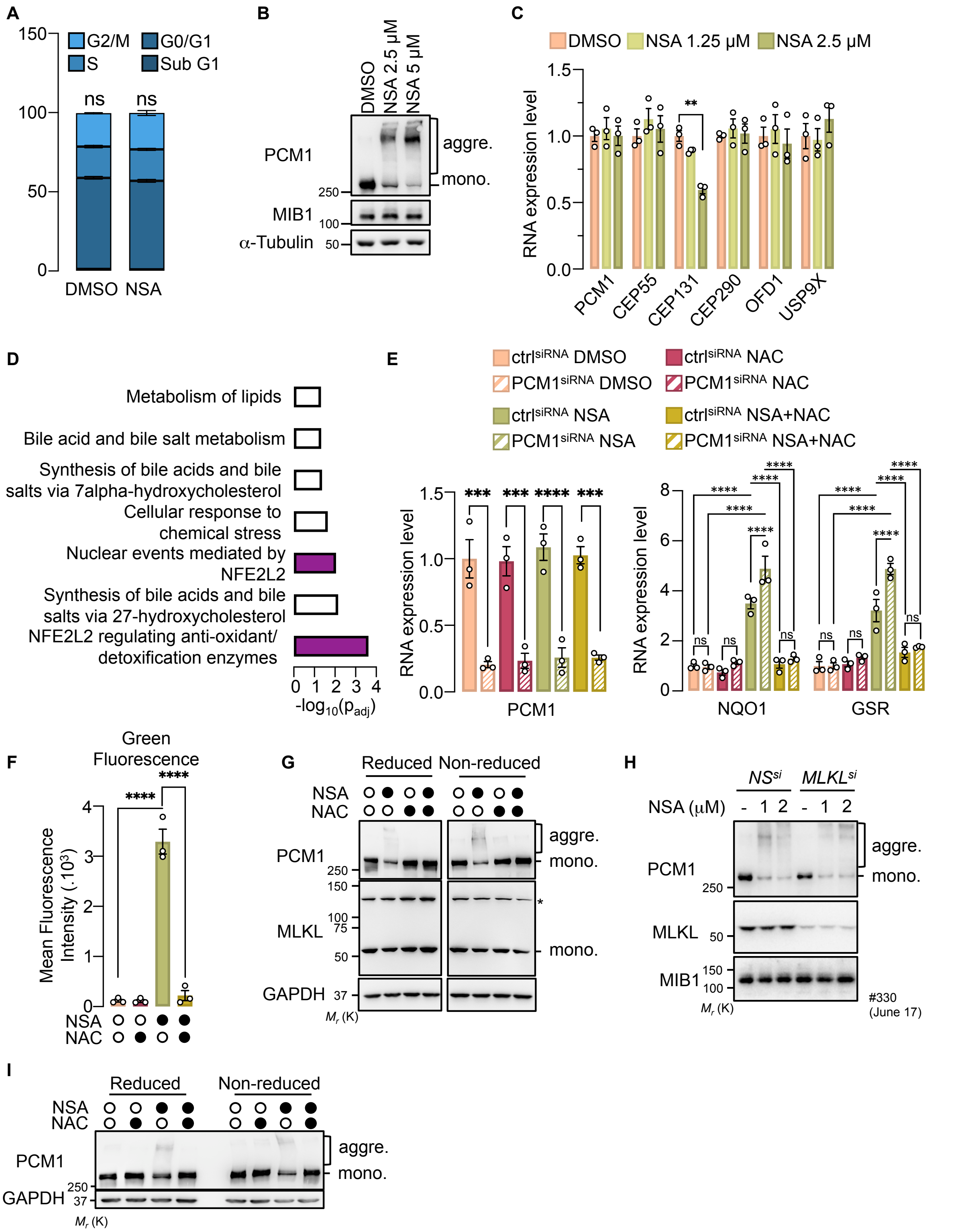


**Figure S1. Necrosulfonamide drives PCM1 aggregation, related to Fig. 1**

(**A**) Cell cycle distribution in Jurkat cells treated with overnight DMSO or NSA (2.5 μM). Histograms represent the mean ± SEM (n=3; ns, non-significant, ANOVA).

(**B**) Cell lysates from RPE-1 cells treated overnight with NSA (2.5 and 5 μM) were prepared and analyzed by Western blotting with antibodies specific to the indicated proteins. Aggre., aggregates; mono., monomeric. Molecular weight markers (*Mr*) are indicated.

(**C**) mRNA analysis by RT-qPCR of Jurkat cells treated with NSA (1.25 or 2.5 μM). Data are presented as the mean ± SEM fold change on three independent experiments using ACTB and HPRT1 as housekeeping genes for normalization, (***P* < 0.01, ANOVA).

(**D**) REAC enrichment analysis of the top pathways from RNAseq transcriptomic analysis of Jurkat cells treated with DMSO or 2.5 μM NSA (shrunken Log2 fold change of at least 1.2; 235 differentially expressed genes).

(**E**) Jurkat cells were transfected with small interfering RNA (siRNA) for PCM1 or scramble non-specific (NS). Cells pre-incubated for 2h with 5 mM N-acetylcysteine (NAC) were treated with 2.5 μM NSA overnight and mRNA analysis by RT-qPCR was performed. Data are presented as the mean ± SEM fold change on three independent experiments using ACTB and HPRT1 as housekeeping genes for normalization, (****P* < 0.001, *****P* < 0.0001, ANOVA).

(**F**) Flow cytometry analysis of green autofluorescence in Jurkat cells treated for 2h with 5 mM NAC before incubation overnight with 2.5 μM NSA (mean ± SEM; n=3; *****P* < 0.0001; ANOVA).

(**G**) Jurkat cells were pretreated with 5 mM NAC for 2h before overnight incubation with NSA, as indicated. Cell lysates were analyzed by Western blotting in reducing (10% 2β-ME) and non-reducing conditions. * shows a non-specific band.

(**H**) Jurkat cells were transfected with small interfering RNA (siRNA) for MLKL or scramble non-specific (NS). Samples were treated and analyzed as in (B).

(**I**) MEFs were treated and analyzed as in (G).

All presented data are representative of three independent experiments.

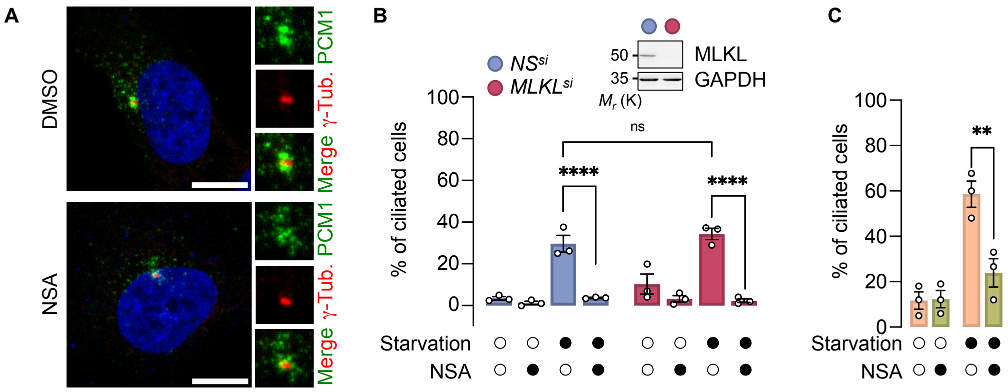


**Figure S2. Necrosulfonamide impacts ciliogenesis independently of MLKL, related to Fig. 2**

(**A**) Confocal microscopy analysis of RPE-1 cells treated overnight with 2.5 µM NSA showing the localization of PCM1 and γ-Tubulin, with nuclei stained by 4'-6-diamidino-2-phenylindole (DAPI). Scale bar, 10 µm.

(**B**) RPE-1 cells were transfected with small interfering RNA (siRNA) for MLKL or scramble non-specific (NS). Cells were serum-starved and treated with 2.5 µM NSA for 24h. Acetylated tubulin (cilia) and PCM1 were immunostained with specific antibodies, and nuclei were stained by DAPI. The histogram shows the quantification of ciliated cells from (means ± SEM of three independent experiments; n > 100 cells counted per sample; ****$P$ < 0.0001, ANOVA). The inset blot shows the efficiency of MLKL silencing.

(**C**) MEF cells were serum-starved and treated with 1.25 µM NSA for 48h, and analyzed as in (B). The histogram shows the quantification of ciliated cells (means ± SEM of three independent experiments; n > 100 cells counted per sample; **$P$ < 0.01, ANOVA).

All presented data are representative of three independent experiments.

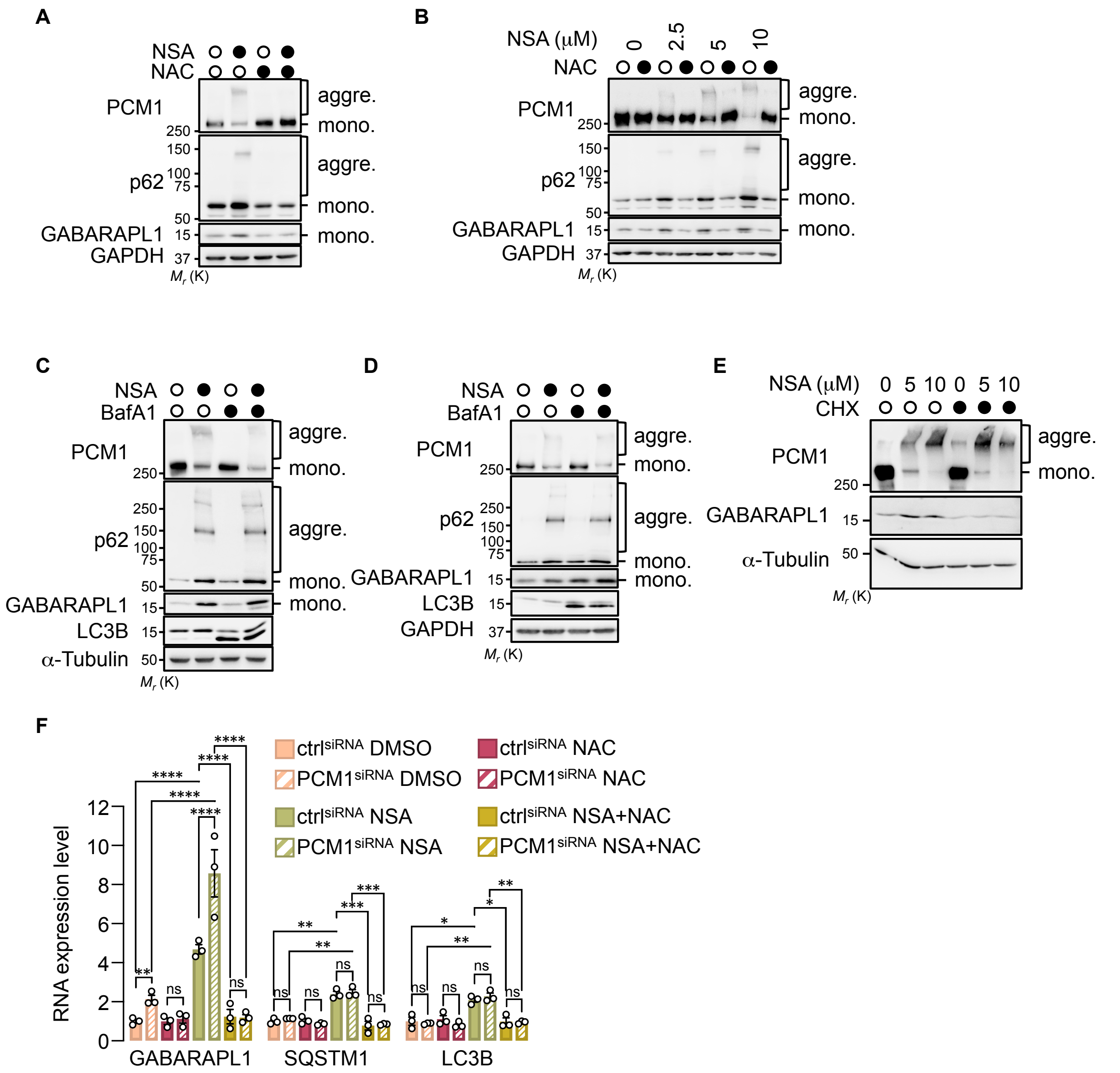


**Figure S3. Necrosulfonamide impairs autophagy in an MLKL-independent Manner, related to Fig. 3**

(**A and B**) L929 cells (A) and MEFs (B) were treated with 5 mM NAC for 2h before overnight incubation with NSA. Cell lysates were analyzed by Western blotting in non-reducing conditions. Aggre., aggregates; mono., monomeric. Molecular weight markers (*Mr*) are indicated.

(**C**) Jurkat cells were pretreated with 100 nM Bafilomycin A1 (BafA1) for 1h and incubated with 2.5 μM NSA overnight. Cell lysates were analyzed by Western blotting with antibodies specific to the indicated proteins.

(**D**) RPE-1 cells were treated and analyzed as in (B).

(**E**) Jurkat cells were incubated with cycloheximide (1 μM) for 1h and treated with 5 or 10 μM NSA for 6h. Samples were analyzed as in (A).

(**F**) Jurkat cells were transfected with small interfering RNA (siRNA) for PCM1 or scramble non-specific (NS). Cells pre-incubated for 2h with 5 mM N-acetylcysteine (NAC) were treated with 2.5 μM NSA overnight and mRNA analysis by RT-qPCR was performed. Data are presented as the mean ± SEM fold change on three independent experiments using ACTB and HPRT1 as housekeeping genes for normalization, (*$P < 0.05$, **$P < 0.01$, ***$P < 0.001$, ****$P < 0.0001$, ANOVA).

All presented data are representative of three independent experiments.

### Table S1. List of the Inhibitors used

| Chemical | Concentration |
|---|---|
| Necrostatin-1s | 20 µM |
| z-VAD-fmk | 20 µM |
| Q-VD | 10 µM |
| Birinapant | 5 µM |
| Rapamycin | 50 nM |
| Everolimus | 20 nM |
| Torin | 100 nM |
| Wortmannin | 50 nM |
| BX795 | 1 µM |
| LY294002 | 10 µM |
| Akti 1/2 | 1 µM |
| Nocodazole | 1 µM |
| Taxol | 10 nM |
| Bortezomib | 10 nM |
| Bisindolylmaleimide VIII (BIM VIII) | 500 nM |
| SB203580 | 10 µM |
| BMS-345541 | 1 µM |
| SP600125 | 20 µM |
| Mepazine (MPZ) | 20 µM |
| PF3644022 | 1 µM |
| Trametinib | 1 µM |
| Amlexanox | 20 µM |
| MRT67307 | 2 µM |
| Hydroxyurea | 10 µM |
| Butylated hydroxyanisole (BHA) | 200 µM |
| Ro-3306 | 5 µM |
| Lovastatin | 10 µM |
| Lenalinomide | 10 µM |
| Axitinib | 5 µM |
| AZD7762 | 100 nM |
| Y-27632 | 10 µM |
| Dasatinib | 25 µM |

### Table S2. List of the Primers used

| |
|---|
| ACTBf: GGACTTCGAGCAAGAGATGG |
| ACTBr: AGCACTGTGTTGGCGTACAG |
| HPRT1f : TGACACTGGCAAAACAATGCA |

| |
|---|
| HPRT1r : GGTCCTTTTCACCAGCAAGCT |
| PCM1f: TTGAAGTGTGGAGCGGGAAA |
| PCM1r: GTTGGGCACCCCAATCCATA |
| CEP131f: TGATGCTCTTCGAGGGCAG |
| CEP131r: GGAACTCCGGGCATTGGAT |
| SQSMT1/P62f: TGCCCAGACTACGACTTGTG |
| SQSMT1/P62r: AGTGTCCGTGTTTCACCTTCC |
| GABARAPL1f: CCCTCCCTTGGTTATCATCCA |
| GABARAPL1r ACTCCCACCCCACAAAATCC |
| LC3Bf: GCTCATCAAGATAATTAGAAGGCG |
| LC3Br: CTGGGAGGCATAGACCATGT |
| USP9Xf: AAGTGAAGCATGTCAGCGATT |
| USP9Xr: GCCACACATAGCTCCACCA |
| CEP290f: AGATGCTCACCGAACAAGTAGA |
| CEP290r: ATGAGTCTGTTGAGAAAGGGTTG |
| CEP55f: TGAAGAGAAAGACGTATTGAAACAA |
| CEP55r: GCAGTTTGGAGCCACAGTCT |
| OFD1f: TCTTTCCAGAAAGTGGTTTGG |
| OFD1r: GAGACTGGAAGTAGGGTTGATTTT |
| GSRf : CACGAGTGATCCCAAGCCC |
| GSRr : CACGAGTGATCCCAAGCCC |
| NQO1f : GAAGAGCACTGATCGTACTGGC |
| NQO1r : GGATACTGAAAGTTCGCAGGG |
